# Infectious Disease Forecasting via Physics-Informed Machine Learning

**DOI:** 10.64898/2026.06.12.731957

**Authors:** Joseph Hart, Hudson Smith, Lior Rennert, Christopher S. McMahan

## Abstract

Infectious disease transmission evolves as a dynamic process shaped by biological mechanisms, population behavior, and intervention policies, yet public health responses are often driven by lagging indicators. Accurate short- and long-term disease forecasting is essential for the timely deployment of intervention strategies, healthcare capacity planning, and uncertainty-aware, risk-informed decision-making. To address this challenge, three broad classes of forecasting models have traditionally been used: statistical, machine learning, and mechanistic approaches. However, each of these modeling paradigms faces fundamental limitations. In particular, traditional statistical models often lack the flexibility needed to capture complex disease dynamics, machine learning approaches require large, high-quality data streams, and mechanistic models are notoriously difficult to calibrate. To overcome these challenges, we propose a novel physics-informed machine learning (PIML) framework for forecasting infectious disease dynamics. Our approach simultaneously forecasts new case and hospitalization counts, along with other key epidemiological quantities such as the time-varying reproduction number. This is achieved through the design of a machine learning model and estimation strategy regularized by a system of differential equations that encode disease dynamics of the SIHR model, thereby bridging the gap between purely data-driven and mechanistic models. We demonstrate the proposed methodology through in-depth numerical studies and an application to COVID-19 data collected in the state of South Carolina.

## 1 Introduction

Recent years have underscored the persistent threat posed by both emerging and endemic infectious diseases. The COVID-19 pandemic demonstrated the profound societal, economic, and healthcare con-sequences of rapidly spreading novel pathogens [1]. At the same time, seasonal respiratory pathogens such as influenza and respiratory syncytial virus (RSV) continue to place substantial recurring burdens on healthcare systems worldwide, contributing to significant morbidity, mortality, and hospital overcrowding [2]. Collectively, these challenges emphasize the critical need for accurate epidemiological forecasting tools capable of informing public health decision-making and resource allocation by enabling proactive responses to emerging infectious disease threats [3]. In particular, accurate forecasts of cases, hospitalizations, and transmission intensity (e.g., time-varying reproductive numbers) allow policymakers and health systems to anticipate surges, allocate essential resources such as hospital beds, staffing, and medical supplies, and implement timely interventions (e.g., vaccination campaigns, social distancing measures, or travel restrictions) [4, 5, 6]. In the absence of forecasting, responses are often delayed and less effective, leading to preventable morbidity, mortality, and economic disruption. Beyond immediate response, infectious disease forecasting plays a role in strategic planning and risk assessment. Forecasting supports the identification of high-risk regions and populations [7], enabling targeted surveillance [8] and intervention strategies [9] that improve efficiency and equity in public health efforts. Forecasting models also provide a framework for evaluating “what-if” scenarios, allowing decision-makers to assess the potential impact of alternative policies before implementation [10, 11, 12].

A wide range of methodologies have been developed for infectious disease forecasting, each broadly falling into one of three categories: traditional statistical approaches, machine learning methods, and mechanistic models. Examples of traditional statistical methods include generalized linear models [13], generalized additive models [14], hierarchical Bayesian models [15], ARIMA [16], SARIMA [17], and Gaussian process models [18], among many others. Going beyond traditional statistical models, a substantial portion of the recent literature has leveraged modern machine learning techniques to forecast the spread of infectious diseases; e.g., random forests [19], gradient boosting [20], neural networks [21], and long short-term memory (LSTM) networks [22, 23]. Though machine learning approaches provide competitive predictive performance, their limited interpretability and lack of mechanistic structure motivate the use of compartmental models that explicitly represent disease transmission processes [24]. In particular, compartmental models, a sub-class of mechanistic models, have become a cornerstone of epidemiological forecasting and can be formulated in variety of manners; e.g., SIR [25], SEIR [26], and SEIHR [27]. Regretfully, all of the aforementioned forecasting methods suffer from key limitations. In particular, classical statistical models often lack flexibility and fail to capture nonlinear dependencies, whereas machine learning methods, though flexible, are computationally expensive, require large, high-quality datasets that may be unavailable in data-sparse settings, and often extrapolate catastrophically when forecasting. Compartmental models, in contrast, are difficult to calibrate and are therefore often specified under simplifying assumptions (e.g., homogeneous mixing and time-invariant parameters), limiting their ability to represent nonstationary transmission dynamics.

To address these limitations, physics-informed neural networks (PINNs) [28] have recently emerged as a promising framework for infectious disease forecasting that blends the strengths of the aforementioned methodologies. Briefly, PINNs train neural networks subject to soft constraints induced by governing known dynamics or physics, often expressed through systems of differential equations, thereby unifying data-driven learning with mechanistic structure [29]. By regularizing the estimation process in this manner, PINNs effectively constrain the search space to solutions that are consistent with the underlying dynamics, which can improve generalization in settings with sparse or noisy data [30]. In infectious disease applications, these dynamics are typically represented by a system of differential equations arising from compartmental models, allowing established epidemiological principles to be incorporated directly into the estimation procedure. For example, PINNs formulations that leverage compartmental model regularization have been successfully applied to infectious disease forecasting with regard to cases, hospitalizations, and deaths as well as to estimate time-varying transmission parameters [31, 32, 33]. The success of PINNs stems from their ability to integrate mechanistic knowledge with flexible function approximation using neural networks. Despite these advantages, PINNs inherit several limitations from neural networks and introduce additional challenges due to the physics-informed formulation. In particular, the regularized loss function is often highly non-convex and poorly scaled, leading to slow or unstable convergence during training [34, 35]. While recent approaches such as loss splitting [36] and adaptive weight balancing [37] aim to improve optimization, they do not fully resolve these issues. Furthermore, training PINNs can be computationally expensive, causing standard uncertainty quantification techniques, such as bootstrapping, to become intractable. Uncertainty estimation in PINNs remains an open challenge.

To overcome these challenges, we propose a novel physics-informed machine learning (PIML) method for the purposes of infectious disease forecasting that is easy to train, provides estimates of key epidemiological parameters of interest, and allows for uncertainty quantification in a straightforward manner. In particular, our approach is designed to jointly model case and hospitalization counts, and leverages the SIHR [38] model to regularize the loss function. Instead of using neural networks, we approximate all of the unknown functions (i.e., the four compartment trajectories and the time-varying parameters of the SIHR model) in our model with integrated M-splines, also known as I-splines [39]. This choice offers several key advantages. First, our formulation leads to fewer trainable parameters when compared to the neural network models commonly adopted in PINNs while retaining sufficient flexibility. Second, our spline formulation leads to a model fitting strategy that is easy to implement and is computationally efficient. This attribute efficiency allows us to leverage boot-strapping techniques to quantify uncertainty. As a part of our model fitting process, we are able to effectively calibrate the SIHR model to regularize the estimation; i.e., in addition to fitting the model to the data, we are also able to learn the underlying dynamical parameters. Using the calibrated SIHR model we are able to estimate key epidemiological quantities of interest (e.g., time-varying reproductive number) as well as develop forecasting methods that directly obey the learned dynamics.

The remainder of this article is organized as follows. In Section 2 we provide our PIML model and the governing dynamics. In Section 3 we detail our model fitting strategy, forecasting approach, and method for uncertainty quantification. In Section 4, we use simulation to investigate the quality of our PIML method relative to existing PINNs techniques under a variety of infectious disease scenarios. In Section 5 we apply our methods to COVID-19 data collected in the state of South Carolina, and Section 6 concludes with a summary discussion.

## 2 Methods

Following the work of Karniadakis et al. [28], to develop our PIML model we embed epidemiological dynamics via a regularization strategy. That is, under this strategy, the model is estimated based on a loss function of the following form

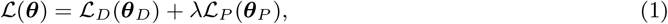

where ***θ***_*D*_ denotes the parameters of the data model, ***θ***_*P*_ denotes the parameters of the governing dynamics, ***θ*** = ***θ***_*D*_ ∪ ***θ***_*P*_ aggregates all of the unknown parameters, with ***θ***_*D*_ ∩ ***θ***_*P*_ ≠ ∅, ℒ_*D*_ (·) represents the data loss, ℒ_*P*_ (·) is the physics loss, and *λ* is the usual penalty parameter. Specifically, ℒ_*D*_ (·) measures the agreement between the model and the observed data, while ℒ_*P*_ (·) measures the agreement between the model and the governing dynamics. This formulation is similar to classical regularization frameworks (e.g., ridge regression and LASSO), where the penalty term shrinks or induces sparsity in the parameter estimates. However, in this context, the regularization term encourages the estimated data model to obey the governing dynamics which are simultaneously calibrated to the observed data. In what follows, we develop both data and physics loss functions that are appropriate for our application.

### 2.1 Data loss

In our application, we jointly model case and hospitalization counts. Let *I*_*t*_(*H*_*t*_) denote the case (hospitalization) counts observed at time *t*, for *t* ∈ **T**_*D*_ = {*t*_1_, …, *t*_*K*_ }. In many physics-informed modeling frameworks for regression, the data loss ℒ_*D*_ (·) is specified to be the mean squared error. However, case and hospitalization counts are typically heteroskedastic, exhibiting a mean-variance relationship in which variability increases with the expected count. Such counts are frequently modeled using Poisson or quasi-Poisson formulations [40, 13] which assume a mean-variance relationship wherein the conditional variance is proportional to the conditional mean, with quasi-Poisson models permitting overdispersion via a dispersion parameter. To capture this structure, we consider a model of the following form for the observed case counts

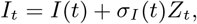

where *I*(*t*) denotes the time-dependent mean function, 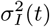 is a time-dependent variance function, and *Z*_*t*_ is a zero mean random variable with unit variance.

Thus, in the absence of the physics based penalty, the unknown mean function may be approximated by a parametric function *I*(*t*; ***θ***_*I*_ ), with estimation based on a weighted least squares criterion of the form

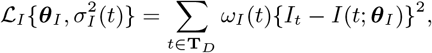

where 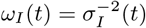. If *I*(*t*; ***θ***_*I*_ ) is linear in ***θ***_*I*_, ***θ***_*I*_ can be efficiently estimated by minimizing 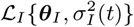 using iteratively reweighted least squares (IRWLS), wherein one alternates between estimating the mean model and updating weights based on the estimated variance function. Using an analogous formulation for the hospitalization counts, we obtain the following joint loss function

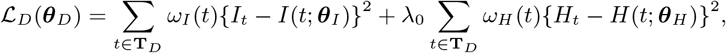

where *H*(*t*; ***θ***_*H*_ ) denotes a parametric model for the time-dependent mean function for hospitalization counts, 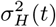 is the corresponding time-dependent variance function, 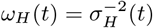, ***θ***_*D*_ aggregates the parameters of the unknown mean and variance functions, and *λ*_0_ controls the relative weighting between the infection and hospitalization data fidelity terms, allowing the model to balance the influence of physics on each data source, which may differ in scale, variability, or measurement reliability. Now that the form of the data loss has been established, we turn attention to specifying the form of *I*(*t*; ***θ***_*I*_ ) and *H*(*t*; ***θ***_*H*_ ).

Since the inputs in our setting consist of a single temporal dimension, we model the two mean functions using spline representations. In particular, we make use of I-splines [39], which leads to the following representation of the mean functions

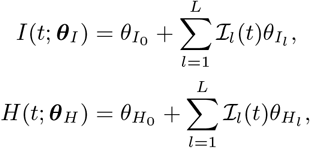

where ℐ_*l*_(*t*), for *l* = 1, …, *L*, are I-spline basis functions, 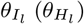 are the corresponding spline coefficients, 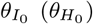 is an intercept term, 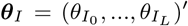, and 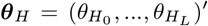. Note that to construct the I-spline basis functions, one needs only to specify the degree (*d*) of the basis splines and choose an increasing sequence of knots (𝒦). Once these specifications are made the I-spline basis functions are fully determined. The degree controls the overall smoothness of the basis functions, while the number and placement of knots determines the overall modeling flexibility, with more knots typically allowing for more flexibility; for further discussion see [39]. In the considered context, the physics loss (derived below) acts analogously to a smoothing loss in penalized splines [41]. This regularizes the estimation of the spline functions thereby mitigating the potential for overfitting. For this reason, we suggest that a moderate number (e.g. 10-30) of equally spaced knots be used with a lower degree spline (e.g., *d* ∈ {1, 2, 3}).

### 2.2. Physics Loss

To formulate our physics loss (i.e., ℒ_*P*_ (***θ***_*P*_ ), we make use of the SIHR model [42], which is an extension of the usual SIR model [43]. Briefly, the SIHR model at any time *t* partitions a population of size *N* into one of four compartments; namely the susceptible *S*(*t*), infected *I*(*t*), hospitalized *H*(*t*), and recovered *R*(*t*) compartments. The temporal flow between these compartments is governed by the following system of ordinary differential equations (ODEs)

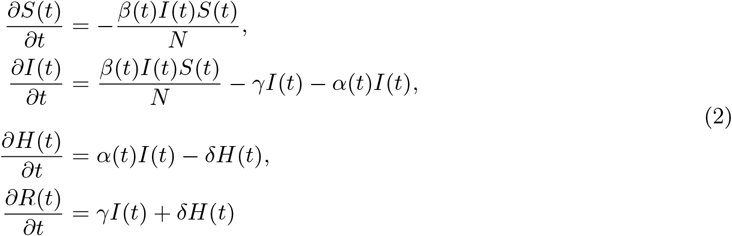

where *β*(*t*) is the transmission rate, *α*(*t*) is the severity (hospitalization) rate, *γ* is the recovery rate, and *δ* is the discharge rate; for further discussion see [38]. A few comments are warranted. First, the transmission and severity rate are often treated as constant, but allowing them to vary over time better reflects real-world conditions [44]. Second, the above system of ODEs assumes no birth or death dynamics, meaning *S*(*t*) + *I*(*t*) + *H*(*t*) + *R*(*t*) = *N*, for all *t* ≥ 0. Third, other key epidemiological parameters of interest can be derived from the quantities above; e.g., the time-varying reproductive number (*R*_*t*_) is given by

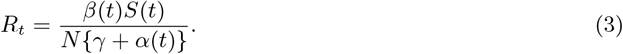

This quantity describes the expected number of secondary infections caused by an infected individual at time *t*, with values of *R*_*t*_ > 1 indicating epidemic growth, *R*_*t*_ = 1 equilibrium, and *R*_*t*_ < 1 decline. Given its interpretability, this parameter is often used to detect changes in disease transmission or asses the effectiveness of interventions [45]. Lastly, given initial values of the compartments and known forms of *β*(*t*), *α*(*t*), *δ*, and *γ* the SIHR trajectories that uniquely satisfy the above system can be found using standard numerical ODE solvers.

Following the squared residual approach [46], we define our physics loss, based on the SIHR model, as follows

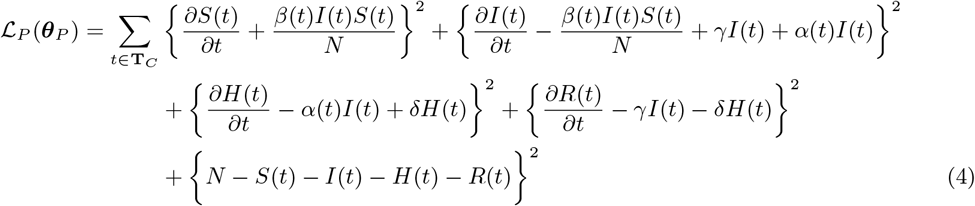

where **T**_*C*_ denotes the set of collocation points at which the physics constraints are enforced. A few comments are warranted. First, as with PINNs, by formulating the physics loss based on the system of ODEs in the manner above, we avoid having to numerically solve the system of ODEs as a part of the model fitting process. Second, the recovery and discharge rates *γ* and *δ* are treated as known constants, which are inferred from the literature [29]. Third, there are several strategies for selecting collocation points (i.e., **T**_*C*_ ), including randomly sampling them at each training iteration or specifying a fixed set, for example **T**_*C*_ = **T**_*D*_, which is the strategy adopted herein. Lastly, and arguebly most importantly, our physics based loss depends on several unknown functions; namely the four compartment trajectories *S*(*t*), *I*(*t*), *H*(*t*), *R*(*t*) and the two time-varying parameters *β*(*t*) and *α*(*t*). Following the strategy outlined in Section 2.1, we approximate these functions using I-splines. The use of I-splines is particularly advantageous in this situation due to the way that they are constructed; i.e., an I-spline basis function of degree *d* is determined as follows

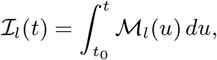

where ℳ_*l*_(*u*) is an M-spline basis function of degree *d* − 1, with both basis functions being defined over the same knot set 𝒦; for further discussion see [39]. This feature provides for a straightforward and analytically tractable way to represent the derivatives appearing in the physics loss; e.g., we have that

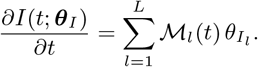

These analytical derivatives eliminate the need for numerical differentiation or autodifferentiation as commonly used in PINNs. This substantially reduces the computational cost associated with evaluating the physics loss.

## 3 Implementation

### 3.1 Model fitting

The goal of our model fitting strategy is to identify ***θ*** ∈ **Θ** that minimizes our proposed loss function; i.e., we seek to identify

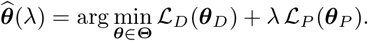

To simplify the exposition of the model fitting strategy, and without loss of generality, we assume that the observation times and the collocation points coincide (i.e.,**T**_*C*_ = **T**_*D*_ ) and that a common spline representation is used for all unknown functions. Under these assumptions, the loss functions can be expressed in matrix form, yielding the following representations for the data and physics losses

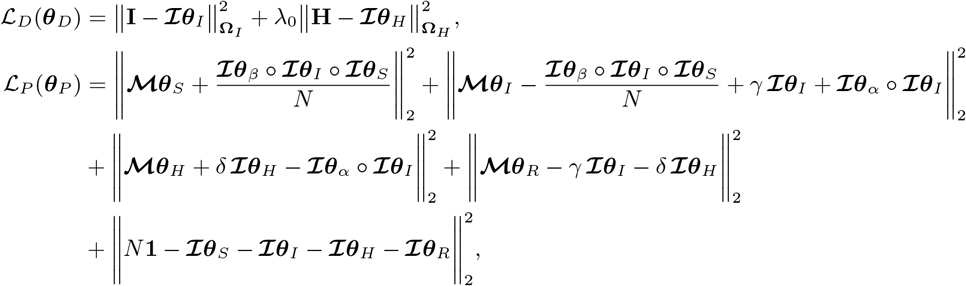

where the vector **I** (**H**) aggregates the observed case (hospitalization) counts, **ℐ** (**ℳ**) is an I-spline (M-spline) design matrix, ***A*** ο ***B*** denotes the Hadamard product between vectors ***A*** and 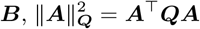 for vector ***A*** and matrix ***Q*, Ω**_*I*_ = diag{*ω*_*I*_ (*t*); *t* ∈ **T**_*D*_ }, **Ω**_*H*_ = diag{*ω*_*H*_ (*t*); *t* ∈ **T**_*D*_ }, 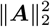 denotes the squared euclidean norm of the vector ***A***, and **1** denotes the all-ones vector. See Appendix A.1 for details on the case when unknown functions have different spline representations and when **T**_*D*_ ≠ **T**_*C*_ .

The above loss admits a structure that enables an efficient model-fitting strategy via block coordinate descent. In particular, holding the variance functions fixed and defining blocks as the spline coefficients associated with each unknown function yields quadratic subproblems that can be solved in closed form.

For example, consider the update for ***θ***_*R*_ which is obtained by solving the following subproblem

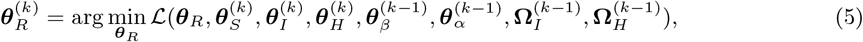

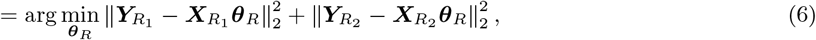

where 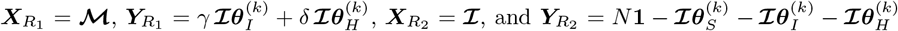. Note, going from (5) to (6) involves dropping the terms in the overall loss function that do not depend on ***θ***_*R*_ and re-expressing the problem under the aforementioned variable declarations. Once this is done, it is straightforward to see that the solution to the stated subproblem is given by 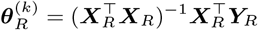 where 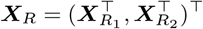 and 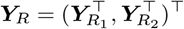. Updates via least squares or weighted least squares solutions for the other sets of spline coefficients follow similarly and the update of the weight matrices (i.e., **Ω**_*I*_ and **Ω**_*H*_ ) are obtained utilizing standard heteroskedastic regression techniques based on the assumed mean-variance structure and the current estimates of ***θ***_*I*_ and ***θ***_*H*_ ; for further discussion see [47] and the strategy outlined in Web Appendix A.1. These steps are repeated in turn until convergence. To initialize our algorithm, experience has shown that using the ordinary least squares estimate of ***θ***_*I*_ and ***θ***_*H*_ aids in speeding convergence. Additionally, to further accelerate convergence, we suggest that *λ* be introduced through a linear schedule increasing from 0 to its target value, analogous to warmstart strategies commonly used in regularized regression where solutions along a regularization path are initialized from previously fitted models. As with other penalized regression methods, the proposed approach relies on aptly specifying *λ* and *λ*_0_. To this end, we make use of standard cross-validation techniques to choose these tuning parameter settings in a data driven manner; for further discussion see Sections 4 and 5. Algorithm 1 succinctly presents our model fitting strategy.

#### Algorithm 1 Model fitting strategy via block coordinate descent

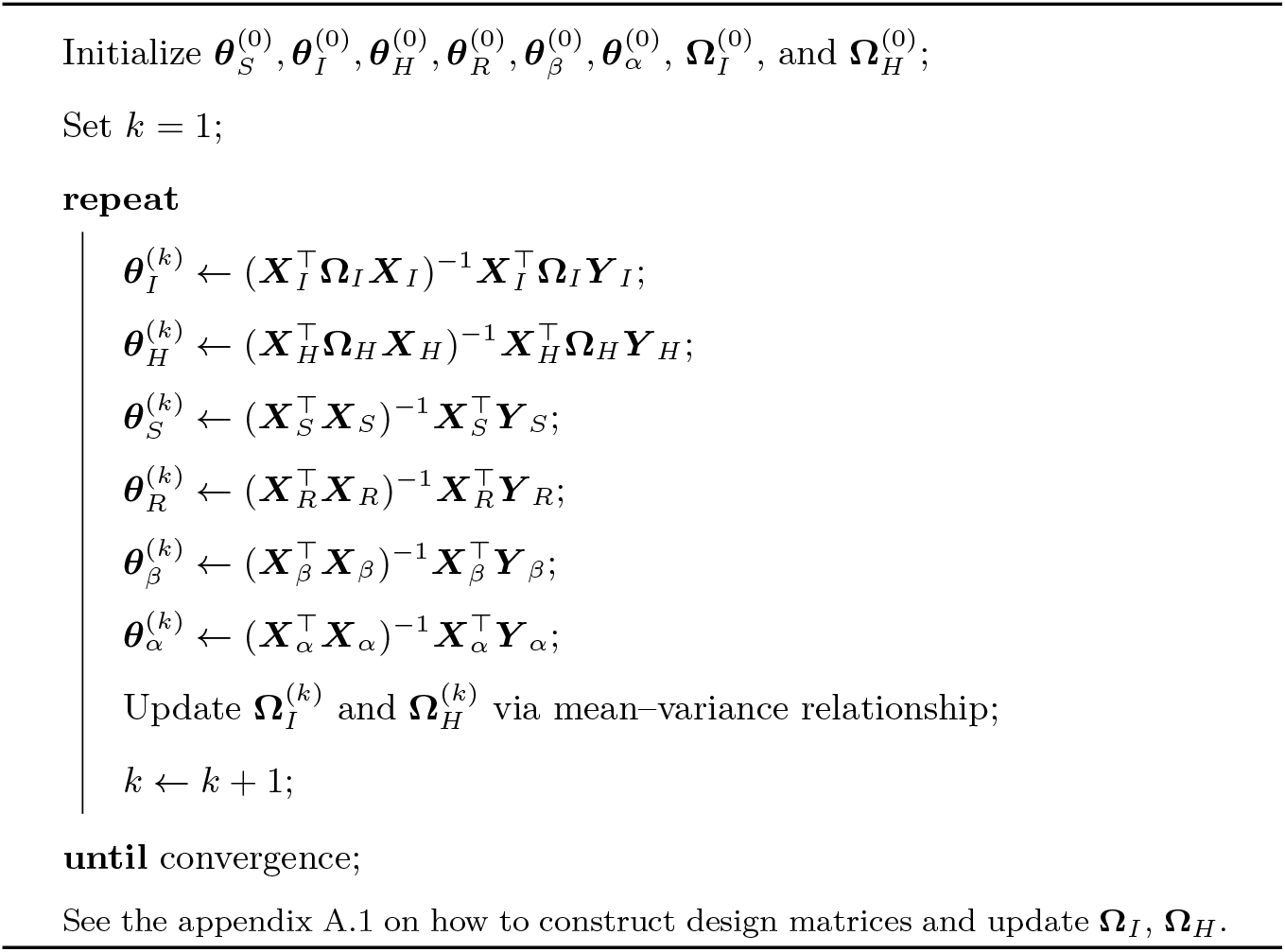

### 3.2 Forecasting and UQ

Many statistical and machine learning epidemiological forecasting methods rely purely on observed data to forecast; they do not leverage underlying disease dynamics. As a result, their forecasts may lack structural consistency with known or inferred epidemiological mechanisms. In contrast, the proposed PIML model uses the SIHR system to impose mechanistic structure on the estimated trajectories. As a by-product of doing this, the proposed approach recovers a data-consistent solution to the SIHR system as a part of the training process; i.e., as a part of the model fitting process described in Section 3.1 we obtain estimates of the compartment trajectories as well as the time-varying parameters, for all *t* ∈ **T**_*D*_ . Of particular interest with regard to forecasting, is the fitted compartment values at the last observed time point (*t*_*K*_ ), which are given by 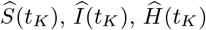, and 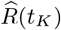. Note, for notational convenience, here and in what follows, we suppress the dependence on ***θ*** in expressing the estimates of the unknown functions; e.g., we write 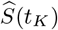 to denote 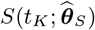, where 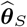 denotes the estimator of ***θ***_*S*_ resulting from our model fitting strategy. These values provide a natural set of initial conditions for forecasting based on the SIHR model. In particular, our strategy to forecasting over a time horizon of length *h* involves solving the SIHR system forward in time using these initial conditions and extrapolated values of the transmission and severity rates. Given that time is our only explanatory variable, we adopt a simple extrapolation strategy that holds the transmission and severity rates fixed at their final fitted values over the forecast horizon; i.e., 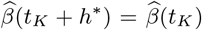 and 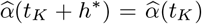, for all *h*^∗^ ∈ (0, *h*). This specification explicitly assumes that the transmission and severity rates are not changing over the forecast horizon, which is reasonable in many scenarios, especially for short forecasting horizons.

Now that our forecasting strategy has been established, we turn our attention to uncertainty quantification. In particular, our goal is to assess the uncertainty associated with the within-sample estimates and to construct prediction intervals for the proposed forecasting procedure. To this end, we employ a bootstrap-based framework for uncertainty quantification. In particular, given the heteroskedasticity commonly observed in epidemiological data, we adopt a wild bootstrap approach [48]. To outline our method, let the fitted trajectories for the case and hospitalization counts be denoted by 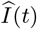 and 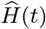, respectively, with corresponding residuals *ϵ*_*I*_ (*t*) and *ϵ*_*H*_ (*t*). Given these quantities, bootstrap training data can be generated in the following manner

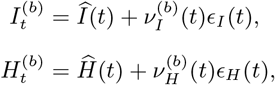

where 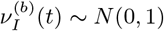 and 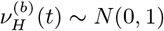. Repeating this process *B* times and training our proposed model on each of the resulting bootstrap data sets yields bootstrap estimates of the four compartment trajectories and the two time-varying parameters; denote these estimates as 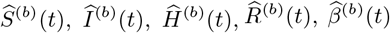, and 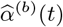, respectively. These bootstrap estimates can be used to draw inference on these unknown quantities as well as functionals (e.g., *R*_*t*_) of these quantities in the usual way; for further discussion see [49] and the strategy outlined in the Appendix A.2.

To provide prediction intervals for our forecast, two sources of variability have to be acknowledged; i.e., the uncertainty associated with estimation of the model parameters and the inherent irreducible error. To aptly account for both of these sources of variability we propose the following strategy. First, following the forecasting method outlined above, we obtain 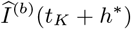 and 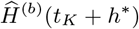, for *h*^∗^ ∈ (0, *h*), by solving the SIHR system of ODEs using 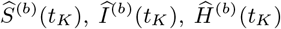, and 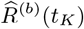 as initial conditions and setting 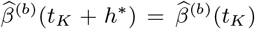 and 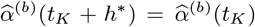. Proceeding in this fashion, we capture uncertainty arising from estimation of the model parameters. To account for the irreducible error in future observations, we employ a second wild-bootstrap step in which pseudo-observations are generated according to

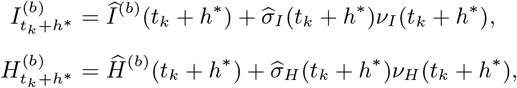

where *ν*_*I*_ (*t*_*k*_ + *h*^∗^) ∼ *N* (0, 1), *ν*_*H*_ (*t*_*k*_ + *h*^∗^) ∼ *N* (0, 1), and 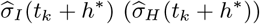 is the time-varying estimated standard deviation of the case (hospitalization) counts. Again, repeating this process B times allows us to create predictions intervals in the usual way; for further discussion see [50]. Algorithm 2 succinctly presents our forecasting and uncertainty quantification strategy. It is worthwhile to point out that bootstrap-based uncertainty quantification can be computationally prohibitive for PINNs because of the considerable computational burden incurred by repeatedly training neural networks across a large number of bootstrap replicates, a limitation that our PIML framework avoids.

#### Algorithm 2 Wild bootstrap for forecast uncertainty quantification

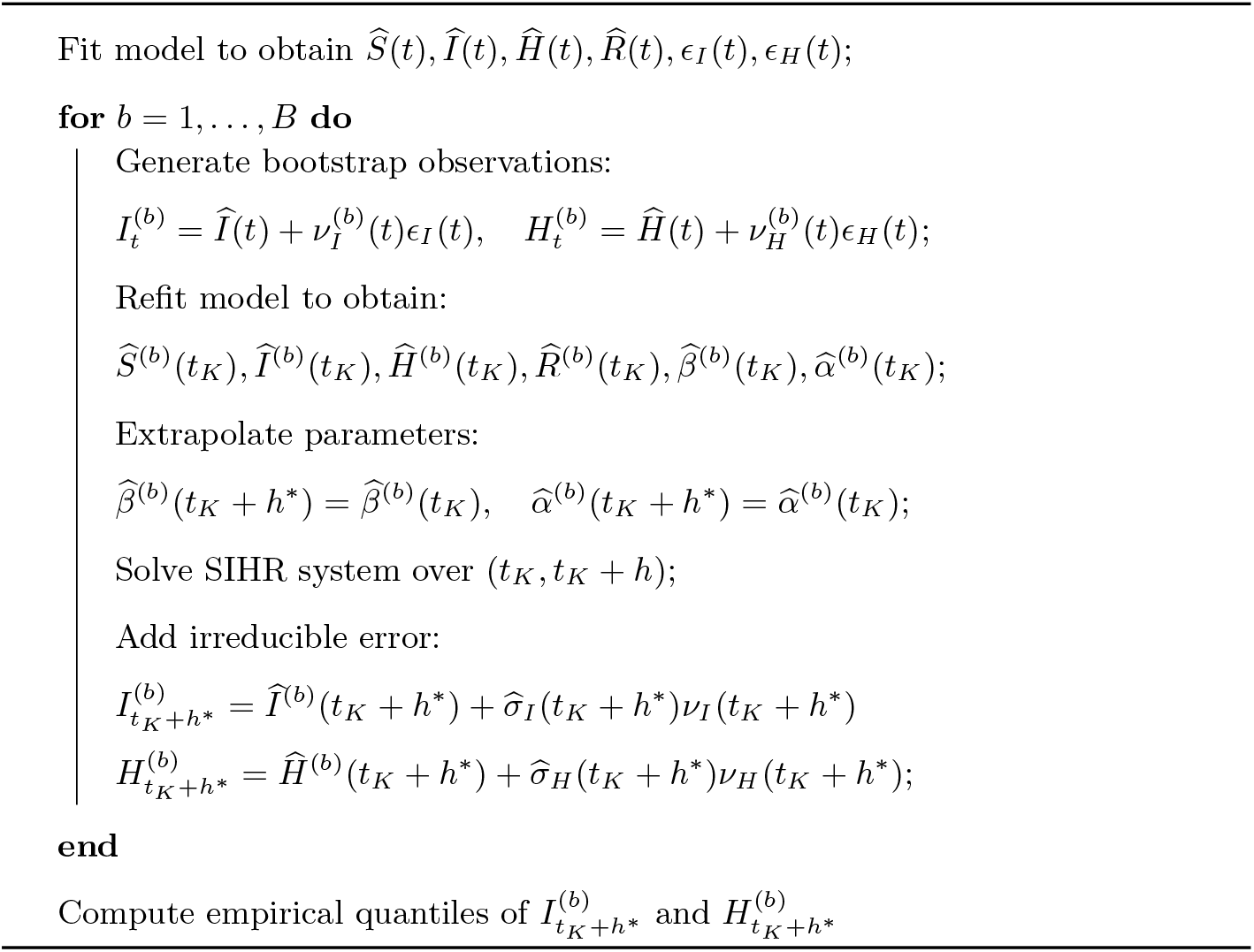

## 4 Numerical Study

To demonstrate the performance of our PIML approach, and to benchmark it against existing PINNs methodologies, we conducted an extensive numerical study. The study was designed to evaluate both the estimation and inferential performance of the proposed method, with emphasis being placed on the within-sample estimation of epidemiological quantities of interest and out-of-sample predictive performance. To this end, we consider two data-generating scenarios: one in which the transmission and severity rates are constant and another in which they vary over time. The former represents a stable disease setting, and the latter reflects dynamically evolving transmission and severity patterns driven by changing behaviors, interventions, or policy decisions. For the constant case we set *β*(*t*) = 0.3 and *α*(*t*) = 0.08 and the time-varying case we set

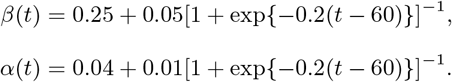

Note, in the time-varying case, these specifications allow for an increase of 0.05 and 0.01 in the transmission and severity rates, respectively, over approximately a 30 day window starting on day 45. For each of these scenarios, we solve the SIHR system of ODEs to obtain *I*(*t*) and *H*(*t*) for a population of size *N* = 500000, with the initial conditions being set to be *S*(0) = 0.98*N, I*(0) = 0.01*N, H*(0) = 0.01*N*, and *R*(0) = 0, and the recovery and discharge rates being given by *γ* = 1/8 and *δ* = 1/9, respectively. These settings were selected to mimic the features of the motivating data; see Section 5 for further discussion. Based on these compartmental trajectories, we generate observed case and hospitalization counts as *I*_*t*_ ∼ *N*{*I*(*t*), *ϕ*_*I*_ *I*(*t*)} and *H*_*t*_ ∼ *N*{*H*(*t*), *ϕ*_*H*_ *H*(*t*)}, respectively, for *t* ∈ **T**_*D*_ = {1, 2, …, 100}. We consider two different settings of the dispersion parameters: the first has the same mean-variance relationship as Poisson data and is achieved by setting *ϕ*_*I*_ = *ϕ*_*H*_ = 1 and the second seeks to mimic the mean-variance relationship of the over-dispersed motivating data (see Section 5), and sets *ϕ*_*I*_ = 80 and *ϕ*_*H*_ = 3. This process is repeated 500 times to create 500 independent data sets under each of the aforementioned settings.

The proposed method was used to analyze each of the simulated data sets. To model all of the unknown functions (i.e., the four compartmental trajectories as well as the transmission and severity rates), we make use of I-splines with knots being placed every fifth time point in **T**_*D*_ . After a preliminary study, we found that setting *d* = 3 for the compartmental trajectories and *d* = 0 for the transmission and severity rate provided adequate performance for both in-sample and predictive inference; however tuning over both *d* and the knot set specification would yield better results as is illustrated in Section 5. To specify the physics based penalty, we set the collocation points to coincide with the observation times; i.e., **T**_*C*_ = **T**_*D*_ . Other configurations of the I-spline representations were considered with no appreciable differences; i.e., we found the proposed approach to be relatively robust to to these specifications. The hyperparameter *λ* was tuned over a grid of 20 values, *λ* ∈ {10^−4.5^, 10}, via cross-validation, while *λ*_0_ was fixed at 1. Jointly tuning *λ*_0_ may further improve the proposed approach. To compare our model against an existing technique, we also used the PINN methodology developed by [51] to analyze each of the data sets, under default settings. All numerical experiments were executed on the Palmetto High-Performance Computing Cluster at Clemson University [52]. To ensure a fair comparison of computational performance across all experiments, both the proposed PIML framework and the PINN baseline were trained on the same high-memory node equipped with dual AMD EPYC 9654 (Genoa) processors (96 cores per socket), yielding 192 physical CPU cores and 750 GB of RAM.

### 4.1 In-sample performance

Figure 1 provides a summary of the results obtained by our PIML approach and the PINN method for the data generated under the constant transmission and severity rates at both error configurations. In this figure we provide average estimates of *I*(*t*), *H*(*t*), and *R*_*t*_ obtained by the PIML and PINN techniques, as well as 0.025 and 0.975 point-wise quantiles of the same. From these results, we see that the proposed PIML technique performs well and outperforms the PINN method. In particular, in the less noisy setting (i.e., *ϕ*_*I*_ = *ϕ*_*H*_ = 1), the average estimates obtained by both the PIML and PINN approaches closely track the true underlying trajectories. However, the PINN based estimates exhibit more variability, particularly for the estimates of *R*_*t*_, than the estimates obtained by our approach. Under increased noise (i.e., *ϕ*_*I*_ = 80 and *ϕ*_*H*_ = 3), these differences become more pronounced. While both methods continue to capture the general trends of the state trajectories, the PINN based estimates exhibit greater variability, and instability in the estimates of *R*_*t*_. The PIML model, on the other hand, remains more stable, producing estimates with less variability. To better quantify the differences in variability, Table 1 reports the ratio of the integrated mean squared error (IMSE) for the functional estimates obtained by the PIML and PINN methods. These results reinforce the assertion that our PIML outperforms the PINN method. That is, the IMSE for estimates from our method are often an order of magnitude, if not more, smaller than those obtained by the PINN approach. Further, based on the average/median of the estimated coverage probabilities reported in Table 1, one can ascertain that our approach also provides reliable in-sample inference; i.e., the estimated coverage probabilities attain their nominal level. It is important to note that the PINN technique does not provide uncertainty quantification or inference. Turning away from estimation performance, we also note that the proposed PIML model is far more computationally efficient than the PINN technique. Total time to validate, train, and obtain uncertainty quantification using our approach was approximately 40 minutes on average for each data set. Specifically, on average it took 2.3 seconds to fit our model. This allowed us to tune *λ* via cross validation over a grid of 20 potential values (approximately 46 seconds on average) and to obtain uncertainty quantification via 1000 bootstrap iterates (approximately 38 minutes on average). In contrast, it took approximately 1 hour on average to simply train the PINN model; this burden makes tuning and bootstrap uncertainty quantification for this competing method computationally prohibitive unlike the proposed approach. We note that the computational cost of our approach can be further reduced through parallelization over both the validation grid and bootstrap replicates.

**Figure 1.**
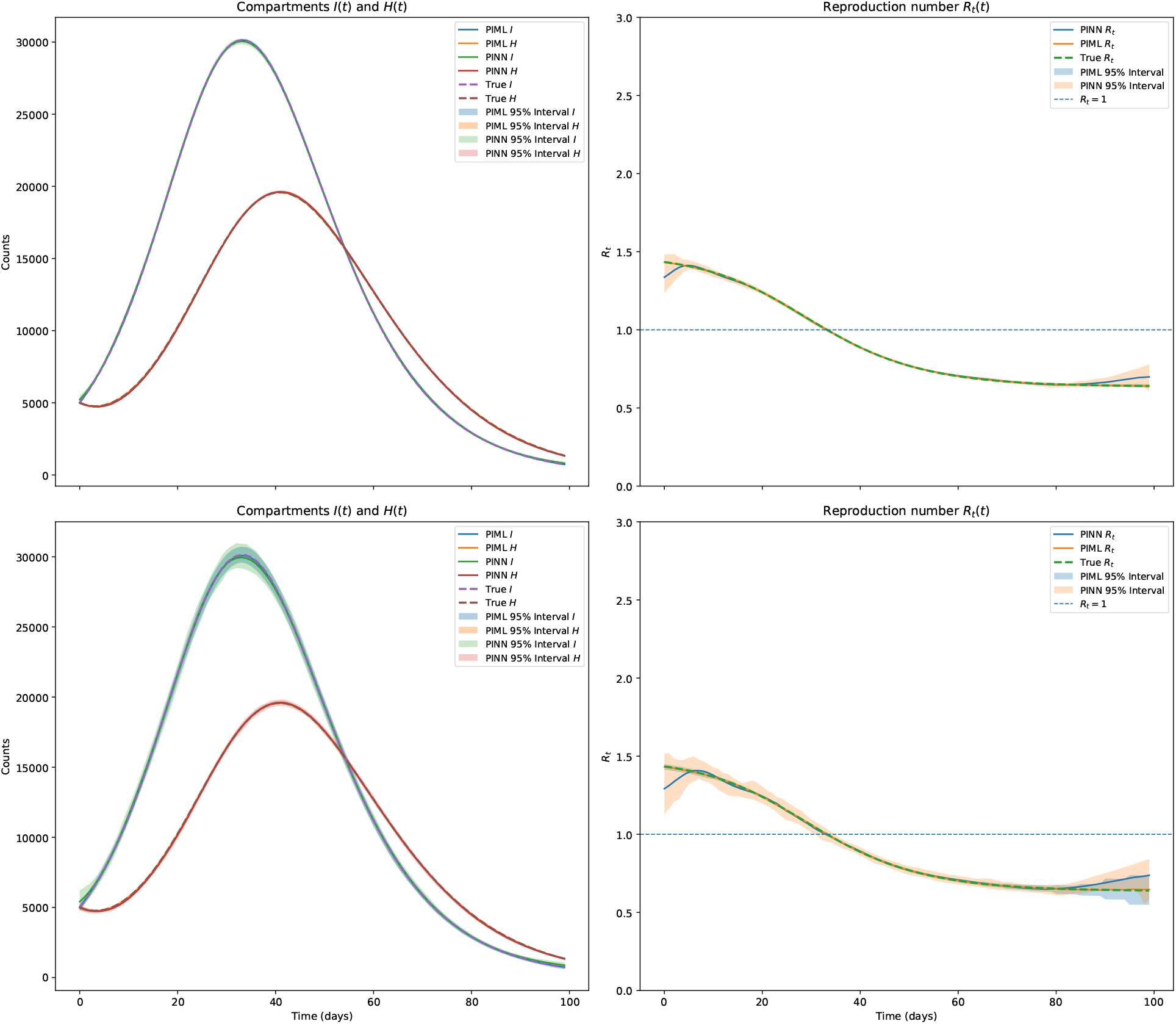
Summary of the estimates obtained by the PIML and PINN methods for the data generated under the constant transmission and severity rates at both error configurations; *ϕ*_*I*_ = *ϕ*_*H*_ = 1 (top row) and *ϕ*_*I*_ = 80 and *ϕ*_*H*_ = 3 (bottom row). Summary of results includes the average estimates of *I*(*t*) and *H*(*t*) (left column), average estimates of *R*_*t*_ (right column), and 0.025 and 0.975 point-wise quantiles of the functional estimates.

**Table 1:**
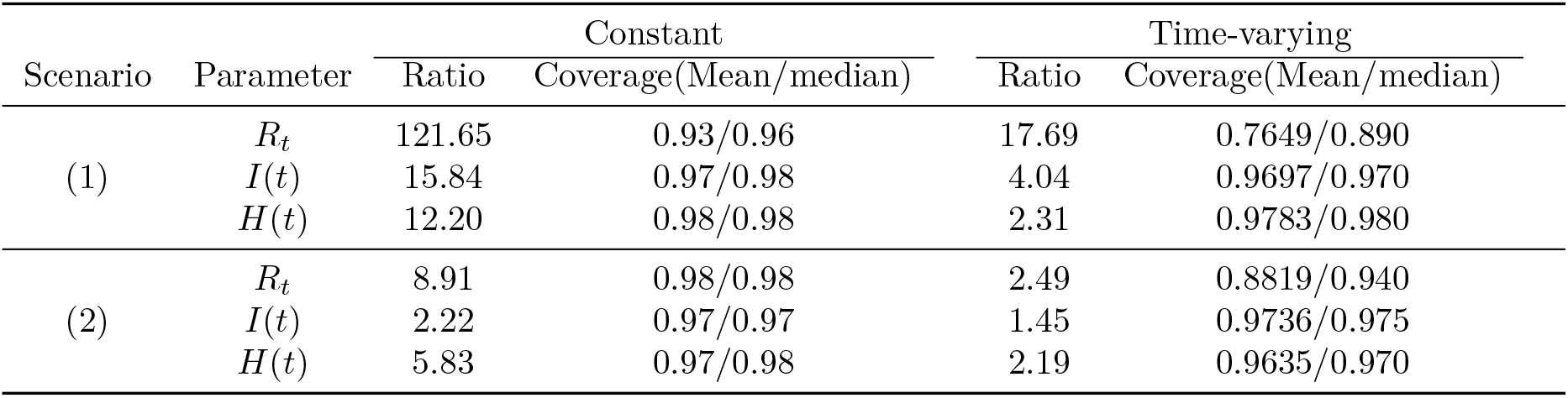
Summary of the results obtained by our PIML approach and the PINN methods when *ϕ*_*I*_ = *ϕ*_*H*_ = 1 (1) and *ϕ*_*I*_ = 80 and *ϕ*_*H*_ = 3 (2), for both constant and time-varying transmission and severity rates. The provided summary reports the ratio of the IMSE for the estimates obtained by the two approaches; i.e., Ratio=IMSE(PINN)/IMSE(PIML). Also provided for our PIML method is the average and median of the estimated coverage probabilities; where averages (medians) are taken over *t* ∈ **T**_*D*_. Note, the PINN method does not provide an approach to quantify uncertainty, thus analogous coverage statistics are not available for this approach.

| Scenario | Parameter | Constant |  | Time-varying |  |
| --- | --- | --- | --- | --- | --- |
|  |  | Ratio | Coverage(Mean/median) | Ratio | Coverage(Mean/median) |
| (1) | $R_t$ | 121.65 | 0.93/0.96 | 17.69 | 0.7649/0.890 |
| | $I(t)$ | 15.84 | 0.97/0.98 | 4.04 | 0.9697/0.970 |
| | $H(t)$ | 12.20 | 0.98/0.98 | 2.31 | 0.9783/0.980 |
| (2) | $R_t$ | 8.91 | 0.98/0.98 | 2.49 | 0.8819/0.940 |
| | $I(t)$ | 2.22 | 0.97/0.97 | 1.45 | 0.9736/0.975 |
| | $H(t)$ | 5.83 | 0.97/0.98 | 2.19 | 0.9635/0.970 |

We now turn to the case in which the transmission and severity rates are time-varying. Figure 2 provides the same summary as depicted in Figure 6, but for the data generating scenario in which the transmission and severity rates are time-varying. Table 1 also provides the IMSE and average estimated coverage probabilities for this setting. These results again reinforce the assertions made above. That is, even when tasked with estimating time-varying transmission and severity rates, our PIML method continues to perform well; i.e., our approach provides efficient and accurate estimates as well as reliable inference. We do, however, observe a slight under coverage associated with the time-varying reproduction number for our approach. Upon further analysis, we found that this occurs in the time range for which the transmission and severity rates are changing; i.e., for coverage probabilities of *R*_*t*_ in *t* ∈ (45, 75). This is attributable to the fact that piecewise-constant splines (i.e., *d* = 0) are being used to approximate both the transmission and severity rate which are continuously varying within this range. Altering our spline specification for these functions and setting *d* = 3 resolves this issue; results not shown. Lastly, our approach continues to outperform the PINN method with regard to estimation performance and at a drastic reduction computational burden. Total time to validate, train, and obtain uncertainty quantification using our approach was approximately 57 minutes on average per dataset (1.2 minutes for tuning and 56 for the bootstrap). In contrast, training the PINN model required approximately 65 minutes on average. Overall, the results of this study tend to suggest that the proposed approach provides both reliable estimates of and inference about key epidemiological parameters.

**Figure 2.**
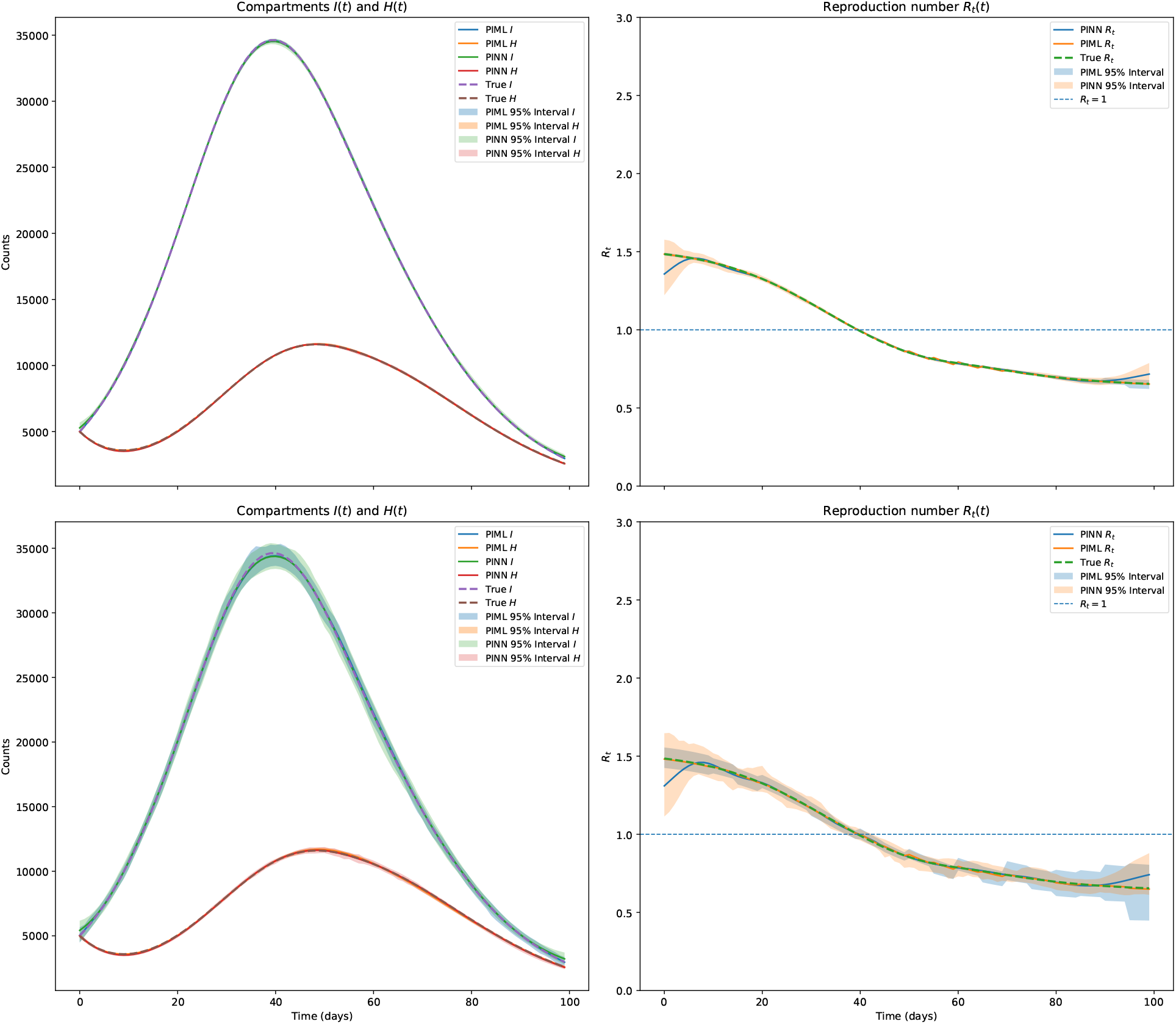
Summary of the estimates obtained by the PIML and PINN methods for the data generated under the time-varying transmission and severity rates at both error configurations; *ϕ*_*I*_ = *ϕ*_*H*_ = 1 (top row) and *ϕ*_*I*_ = 80 and *ϕ*_*H*_ = 3 (bottom row). Summary of results includes the average estimates of *I*(*t*) and *H*(*t*) (left column), average estimates of *R*_*t*_ (right column), and 0.025 and 0.975 point-wise quantiles of the functional estimates.

### 4.2 Forecasting performance

In what follows, we examine the forecasting ability of the proposed PIML technique. In particular, we focus on forecasting three specific epidemiological quantities of interest: case and hospitalizations counts as well as the time-varying reproductive number (i.e., *R*_*t*_). The former two metrics are of primary interest as they directly allow for informed decision making with regard to resource allocation (e.g., hospital beds, staffing, and medical supplies), where as the latter provides insight into the state of the outbreak; e.g., are surges (*R*_*t*_ > 1) or declines (*R*_*t*_ < 1) on the horizon. To explore the performance of the proposed approach in this venue, we make use of the data generating mechanisms described above with *ϕ*_*I*_ = 80 and *ϕ*_*H*_ = 3 (i.e., the more challenging scenario) and generate 500 independent data sets under each setting. Once the data is generated, we consider forecasting the aforementioned quantities over a two-week horizon at different points (*t*_*K*_ ) throughout the surge. In particular, we train our model on the data up until time *t*_*K*_ and then forecast the quantities over *t* ∈ {*t*_*K*_, *t*_*K*+13_}, with *t*_*K*_ ∈ {20, 30, 40, 50}. A few comments are warranted. First, these settings were chosen to challenge the proposed approach with respect to data availability; for example, when *t*_*K*_ = 20, the model is trained using only 20 observations for each of the case and hospitalization count series. Second, the choices of *t*_*K*_ were made so that the forecasting horizon coincides with the peak/decline of the outbreak; time periods that are notoriously hard to forecast. Lastly, in examining the cases in which the transmission and severity rates are both constant and time-varying provides us the opportunity to assess the impact of the forecasting assumption made in Section 3.2; i.e., that transmission and severity rates are constant over the forecasting horizon. For each of these scenarios, we trained our model in exactly the same way as described above, and produced forecasts according to the strategy outlined in Section 3.2. Further, we also implemented the PINN framework outlined in [51] as a competing technique.

Figure 3 summarizes the forecasting results, obtained by both out PIML technique and the PINN approach, for the case in which the transmission and severity rates are constant. In particular, these plots provide the average model fit up to time *t*_*K*_ and the average forecast for *t* ∈ {*t*_*K*_, *t*_*K*+13_}, as well as 0.025 and 0.975 point-wise quantiles of the same. Also provided are point-wise estimated coverage probabilities associated with 95% boot-strap prediction intervals for the forecasted case and hospitalization counts, as well as 95% boot-strap confidence intervals for the time-varying reproductive number. From these results, one will note that the proposed PIML method performs well; i.e., the average forecasts match the true trajectories of the case and hospitalization counts as well as the time varying reproductive number, across all four of settings of *t*_*K*_, and the estimated coverage probabilities for the prediction (confidence) intervals for the forecasted values attain their nominal level. Moreover, our approach is able to accomplish this in incredibly data sparse settings. In contrast, the same cannot be said about the competing PINN method. In particular, the average forecast obtained from this competing technique rarely agrees with the truth, the variability of the forecasted values is far larger, and the PINN method again provides no way to assess uncertainty.

**Figure 3.**
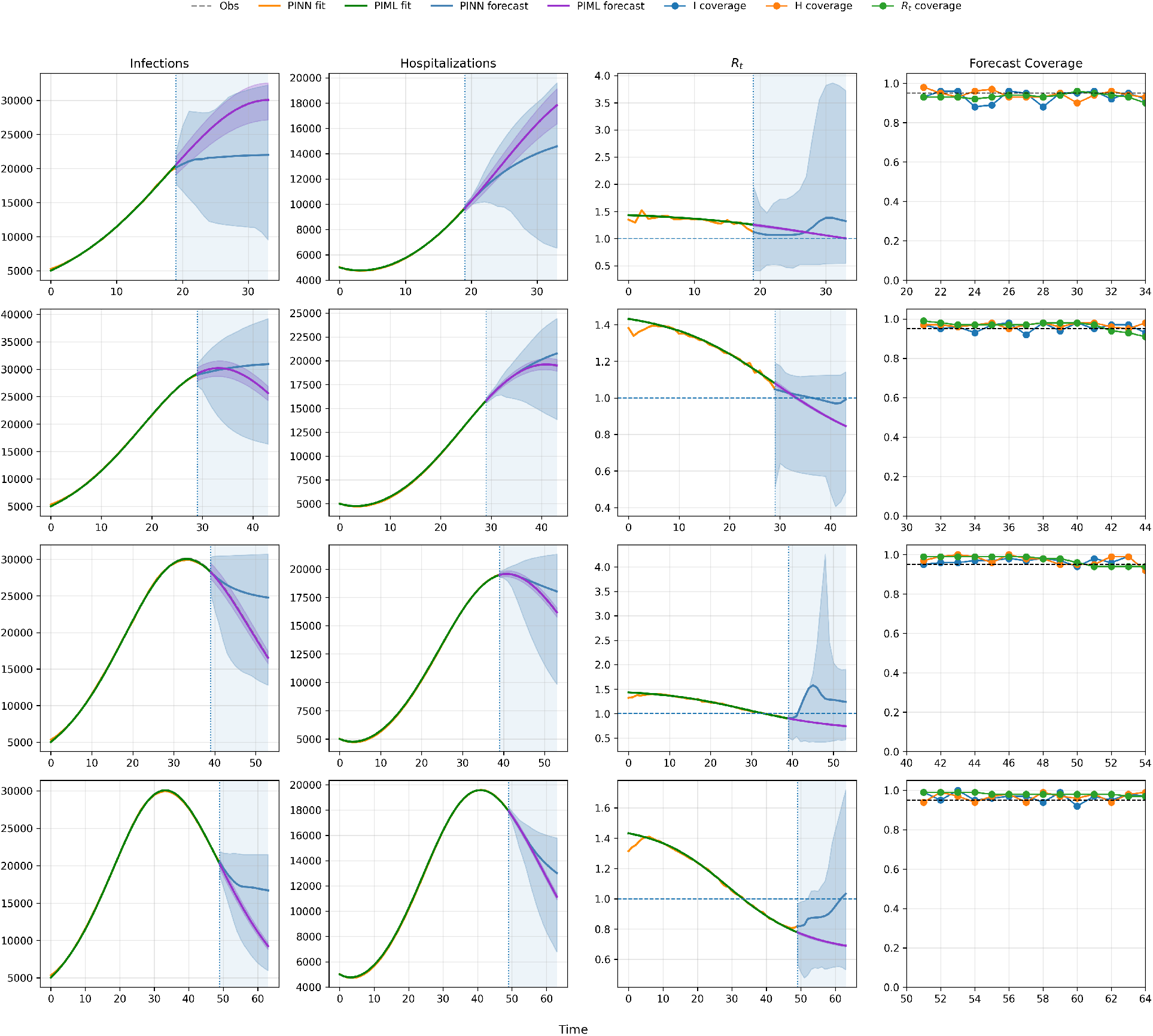
Summary of the estimates obtained by the PIML and PINN methods for the data generated under the constant transmission and severity rates with error configurations *ϕ*_*I*_ = 80 and *ϕ*_*H*_ = 3. Summary of results includes the average estimates of *I*(*t*) and *H*(*t*) (left and second column), average estimates of *R*_*t*_ (third column), and day by day coverage for forecasts (right column. Also included are 0.025 and 0.975 point-wise quantiles of the functional estimates.

Figure 4 provides the same summary as depicted in Figure 3, but for the data generating scenario in which the transmission and severity rates are time-varying. These results again reinforce the assertions made above. That is, even when tasked with estimating time-varying transmission and severity rates, our PIML method continues to perform well; i.e., our approach provides efficient and accurate estimates as well as reliable inference. However, the reliability of the inference does decline when the assumption that the transmission and severity rates are constant throughout the forecasting horizon is violated; e.g., see panels corresponding to *t*_*K*_ = 40 (mild violation) and *t*_*K*_ = 50 (more severe violation). Moreover, our approach continues to outperform the PINN method. In summary, this study has demonstrated that the proposed PIML technique provides reliable forecasts of three specific epidemiological quantities of interest during a challenging forecasting period and based on limited data. Given the accuracy of these forecasts, other quantities of interest could be reliably obtained; e.g., one could predict the timing and duration of outbreak peaks, when and how long hospitalization surges will persist, or when an outbreak will surge (*R*_*t*_ > 1) or decline (*R*_*t*_ < 1).

**Figure 4.**
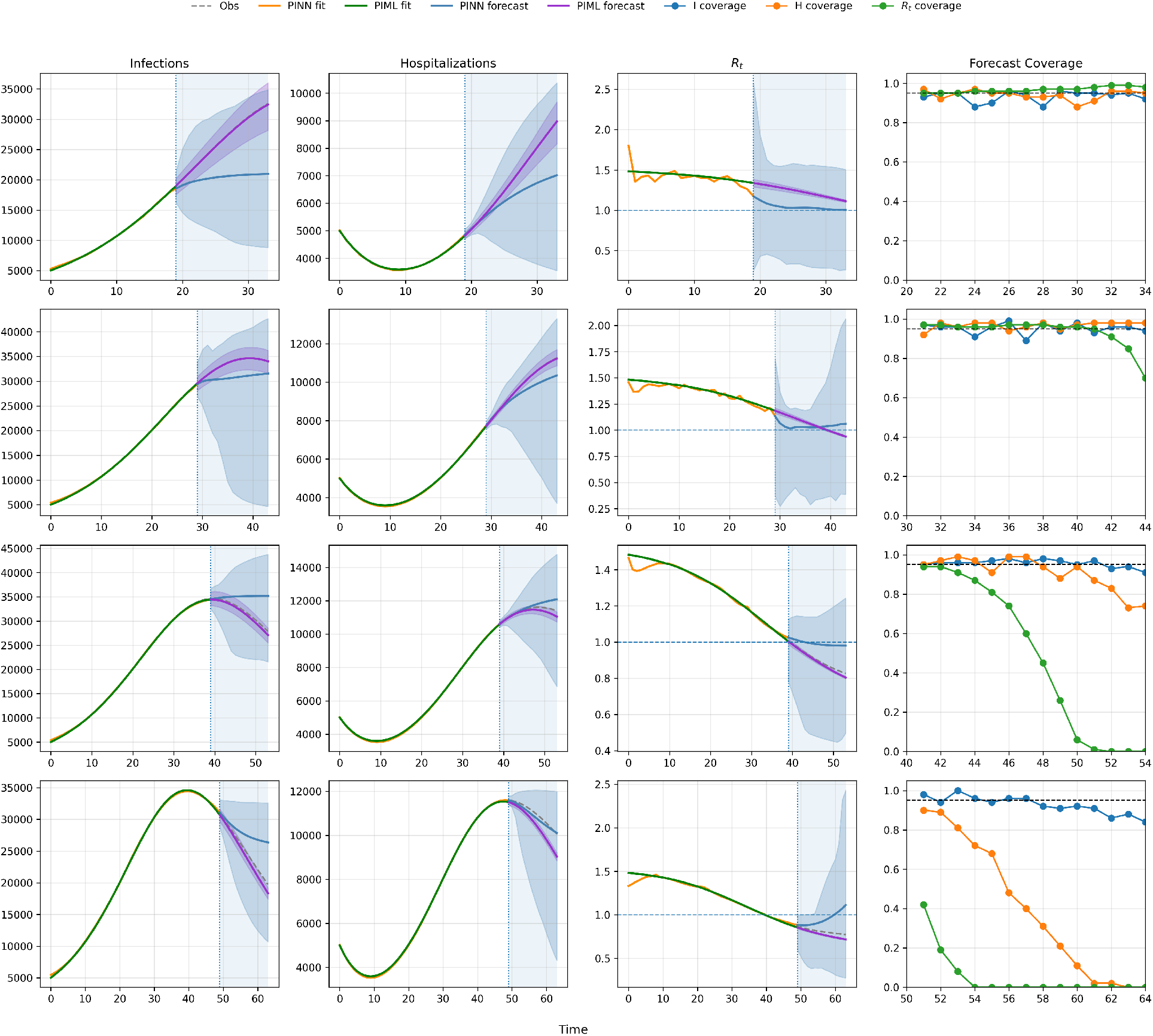
Summary of the estimates obtained by the PIML and PINN methods for the data generated under the non-constant transmission and severity rates with error configurations *ϕ*_*I*_ = 80 and *ϕ*_*H*_ = 3. Summary of results includes the average estimates of *I*(*t*) and *H*(*t*) (left and second column), average estimates of *R*_*t*_ (third column), and day by day coverage for forecasts (right column. Also included are 0.025 and 0.975 point-wise quantiles of the functional estimates.

## 5 Data Application

We further demonstrate the performance of the proposed PIML methodology using COVID-19 data collected from January 1, 2021 to November 1, 2021 in South Carolina. This period represents a challenging forecasting period for a variety of reasons. First, during this timeline the Alpha variant (B.1.1.7), which was more transmissible [53] and severe [54] than the original lineage, began appearing in the United States, quickly becoming the dominant strain; e.g., the Medical University of South Carolina (MUSC) reported that by June 2021 that 78.4% of sequenced cases were the Alpha variant [55]. Second, the Pfizer, Moderna, and Johnson & Johnson vaccines gained emergency use authorization during this period, with broad availability to adults and children over the age of 12 achieved by May 2021 [56]. The initial portion of this period captures a surge in cases likely attributable to increased mobility and indoor social gatherings associated with the holiday season, compounded by winter weather conditions that favored indoor transmission [57]. After a decline in cases, beginning in late summer, the United States experienced a substantial resurgence associated with the emergence and rapid dominance of the Delta variant [58]. This variant was again not only more transmissible than previous strains, but also more severe, exhibiting a higher risk of hospitalization among infected individuals [59]. The confluence of these various factors makes for a challenging forecasting period, during which the transmission and severity rates are expected to be varying.

The data considered for this analysis consists of new daily infection and hospitalization counts, with the former being collected by the New York Times [60] and the latter by the South Carolina Revenue and Fiscal Affairs Office (South Carolina Revenue and Fiscal Affairs Office, 2025) [61]. In this analysis, we focus on assessing our PIML model’s ability to forecast three specific epidemiological quantities of interest: case and hospitalizations counts as well as the time-varying reproductive number (i.e., *R*_*t*_). In particular, we forecast each of these quantities over a two-week horizon at *t*_*K*_ ∈ {100,, 200}, where the data before *t*_*K*_ was used as a training set. To train the model, we set the mean recovery and discharge periods to be *γ* = 1/8 days and *δ* = 1/9 days, consistent with reported values by the Centers for Disease Control and Prevention (CDC) [62]. To ensure consistency between the observed data and these assumptions, raw daily case and hospitalization counts were transformed into estimates of the total infectious and hospitalized populations using 8-day and 9-day rolling sums, respectively. Due to the computational efficiency of our PIML approach, in this study, we consider tuning the penalty parameter *λ*, the parameter that controls the relative weighting between the infection and hospitalization data fidelity terms *λ*_0_, and the spline representations used to model the unknown functions. To this end, we consider uniformly spaced grids of 10 candidate values for both *λ* and *λ*_0_, with *λ* ∈ {0.01,, 0.05} and *λ*_0_ ∈ {0.5,, 1.5}. The unknown compartmental trajectories were modeled using cubic (*d* = 3) I-splines, while the transmission and severity rates were represented using piecewise-constant (*d* = 0) I-splines, with knots being placed at regular intervals of *q*_*c*_ and *q*_*r*_, respectively, with with *q*_*c*_ ∈ {8, 10,, 24} and *q*_*r*_ ∈ {3, …, 6}. To specify the physics based penalty, we again set **T**_*C*_ = **T**_*D*_ . Selection of the spline and regularization tuning parameters was carried out via forecasting-based cross-validation; i.e., for a forecast origin at *t*_*K*_, models are fit using data observed through *t*_*K*−14_, and predictive performance is evaluated over the holdout period [*t*_*K*−13_, *t*_*K*_ ]. The tuning configuration minimizing the validation error was subsequently used to fit the final model. This exploration required fitting our model 3600 times for validation, once to obtain the final model fit, and *B* = 1000 times to drive inference via our bootstrap strategy, resulting in 464,701 total fits across the 101 forecasting windows; which was accomplished, after parallelization across 100 CPUs, in less than 6 hours or approximately 4.5 seconds per model fit.

Figure 5 presents representative model fits, forecasts, and associated confidence and prediction intervals obtained at four forecasting origins (*t*_*K*_ ∈ 140, 160, 180, 200) for COVID-19 case counts, hospitalization counts, and the time-varying reproduction number, (*R*_*t*_). For comparison, corresponding fits and forecasts from the PINN approach are also displayed. A comprehensive summary of forecasting performance across all 101 forecasting windows is provided in Table 2, which reports the correlation, similarity, RMSE, and MAE metrics for both case and hospitalization forecasts. Collectively, these results demonstrate that the proposed PIML framework provides accurate forecasts and consistently outperforms the competing PINN approach across the considered evaluation metrics. In particular, the proposed method successfully captures the highly dynamic nature of COVID-19 transmission during this challenging period, as evidenced by the substantial temporal variability observed in the estimated reproduction number. Moreover, Figure 6 illustrates that the proposed framework yields reliable uncertainty quantification, with prediction intervals attaining their nominal level when the assumption in Section 3.2 holds; i.e., that transmission and severity rates are constant over the forecasting horizon. In contrast, we found that the PINN based forecasts, and model fits, often were drastically more variable and exhibited a greater degree of bias. Finally, we also note that through tuning *λ, λ*_0_, and the I-spline representations, that we found the performance of the proposed approach was relatively robust to the specification of these quantities; i.e., the overall performance for many configurations of these parameters exhibited no appreciable differences. The ability to accurately forecast case counts and hospitalizations, while simultaneously quantifying the associated uncertainty, during such a turbulent period can facilitate proactive resource planning and allocation, enabling healthcare systems to better manage hospital capacity, staffing requirements, and medical supply inventories.

**Figure 5.**
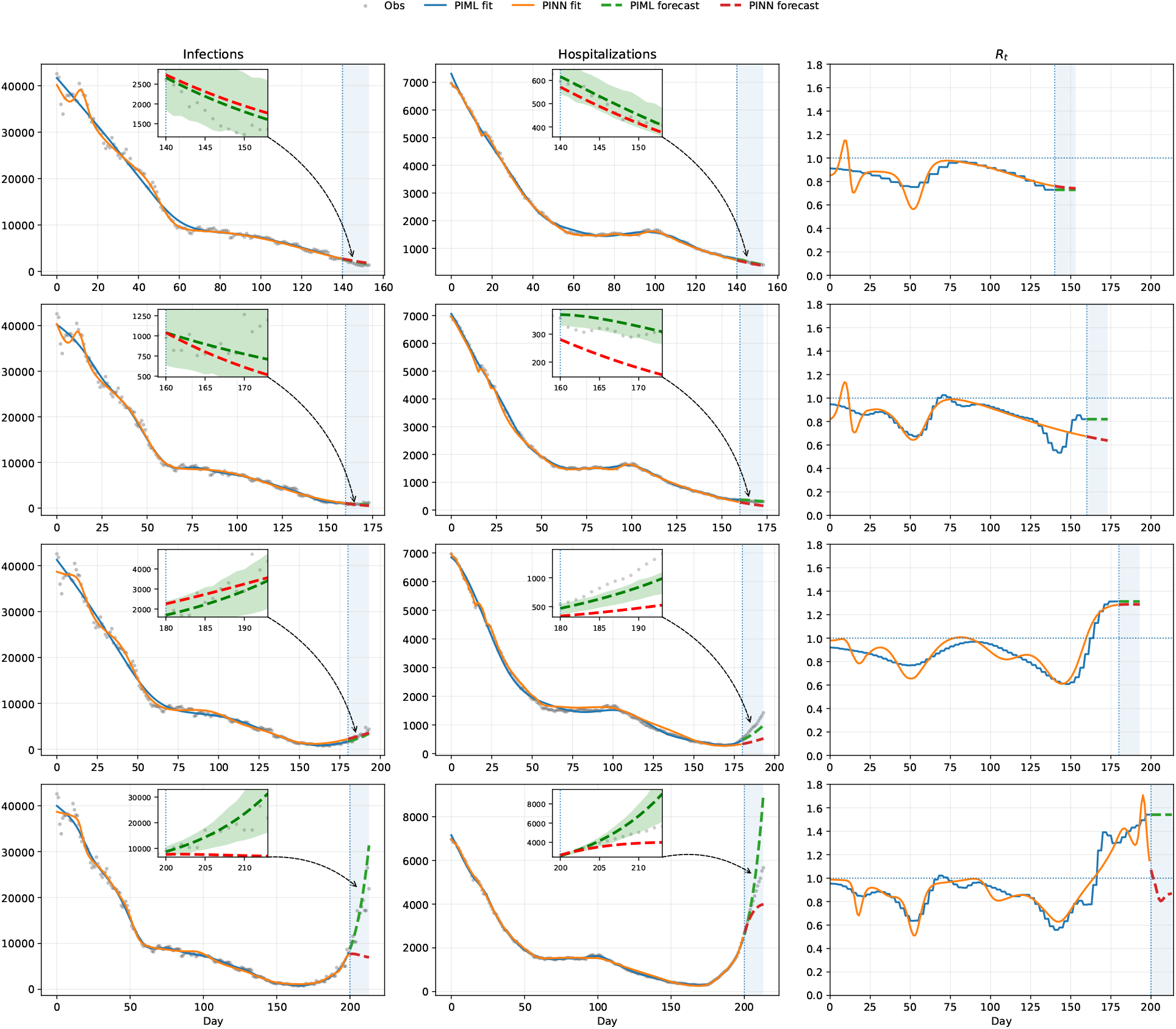
Summary of the model fits, forecasts, and associated confidence and prediction intervals obtained at four forecasting origins for COVID-19 case counts (left column), hospitalization counts (center column), and *R*_*t*_ (right column) by our proposed PIML approach. Also provided are the model fits and forecasts obtained from the competing PINN method. Rows 1-4 correspond to the forecasting origins *t*_*K*_ ∈ 140, 160, 180, 200, respectively. The 14 day forecasting horizons are demarcated by the blue shaded regions in each panel.

**Figure 6.**
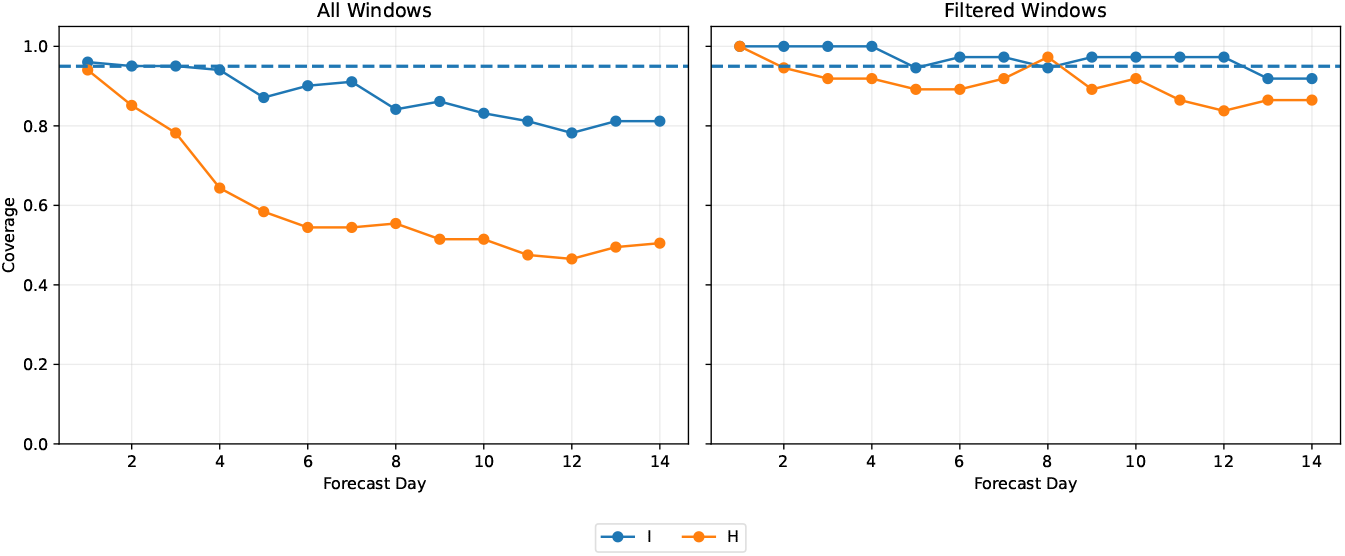
Presented results include the estimated coverage probabilities for the prediction intervals obtained by the proposed PIML approach stratified by day. The panel on the left summarizes the estimated coverages for all 101 14-day forecasts, while the panel on the right summarizes the estimated coverages for the case where the assumption that the transmission and severity rates are constant over the forecasting horizon holds.

**Table 2:** Summary of forecasting performance across all 101 14-day forecasting windows for the proposed PIML approach and the competing PINN method. The summary provides the average correlation, similarity, relative RMSE, and relative MAE between the forecast and the observed data. The relative RMSE and MAE are defined as 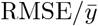 and 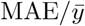, respectively, where 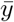 is the mean of the observed values within each forecasting window, where as the similarity is defined in [63].

| Metric | PIS |  | PINN |  |
| --- | --- | --- | --- | --- |
| | $I(t)$ | $H(t)$ | $I(t)$ | $H(t)$ |
| Correlation | 0.665 | 0.831 | 0.524 | 0.659 |
| Similarity | 0.834 | 0.871 | 0.758 | 0.809 |
| Relative RMSE | 0.226 | 0.168 | 0.335 | 0.236 |
| Relative MAE | 0.188 | 0.142 | 0.290 | 0.205 |

Beyond forecasting case counts and hospitalizations, the proposed framework yields epidemiologically meaningful quantities that can directly inform public health decision-making. In particular, the estimated time-varying reproduction number, *R*_*t*_, can serve as an early warning indicator of changes in transmission dynamics before such changes become evident in the observed surveillance data. For example, Figure 5 shows that when *t*_*K*_ = 160, the estimated reproduction number begins to increase around day 145, despite there being little indication of an impending surge in either the case counts or hospitalization data. Although the estimated *R*_*t*_ remains below one during this period, the sustained upward trend suggests that transmission is intensifying and that the epidemic may be approaching a critical threshold. Indeed, *R*_*t*_ subsequently exceeds one, leading to a surge in infections that becomes apparent around day 165. Notably, evidence of this surge does not become visible in the observed case and hospitalization counts until approximately days 165 and 175, respectively. Consequently, the estimated increase in *R*_*t*_ provides policymakers with nearly a month of advance warning, creating an opportunity to proactively implement mitigation strategies, allocate healthcare resources, and prepare for increased healthcare demand before the surge is reflected in traditional surveillance metrics.

## 6 Discussion

In this work, we developed a novel physics-informed machine learning framework for infectious disease forecasting that combines the flexibility of modern statistical/machine learning methods with the interpretability of mechanistic epidemiological models. By leveraging the SIHR model as a regularizing component within the loss function and representing all unknown functions using I-spline basis expansions, the proposed approach avoids many of the computational and optimization challenges commonly encountered with physics-informed neural networks. The resulting methodology is straightforward to train, computationally efficient, and capable of jointly modeling case and hospitalization data while simultaneously learning the underlying disease transmission dynamics. Moreover, the spline-based formulation substantially reduces the number of trainable parameters, facilitating stable estimation, computationally efficient model tuning, and enabling bootstrap-based uncertainty quantification. Beyond generating accurate forecasts, the proposed framework provides estimates of key epidemiological quantities, including the time-varying reproduction number, thereby yielding insights into the underlying state of an outbreak. The performance of the proposed PIML framework was assessed through extensive numerical studies and was further validated through an application to COVID-19 case and hospitalization data collected in South Carolina during 2021. To further disseminate this work, we have developed code, written in Python, that implements all aspects of this work and we have made if freely available in the following github repository.

**Web Appendix**

## A Supplemental Information

### A.1 Additional Details on Block Coordinate Descent Updates

For completeness, we explicitly state the matrix quantities appearing in Algorithm 1. The update for ***θ***_*S*_ is obtained by the closed form update 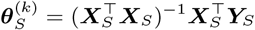, where

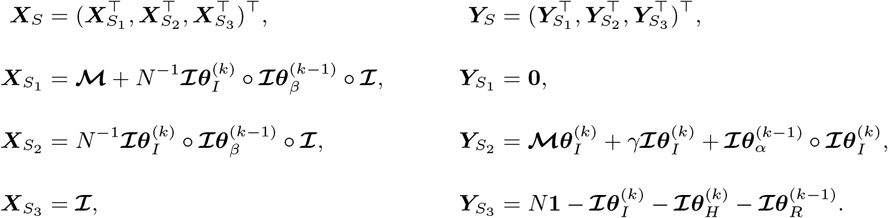

Similarly, the update for ***θ***_*I*_ is given by 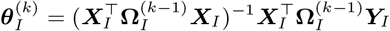, where

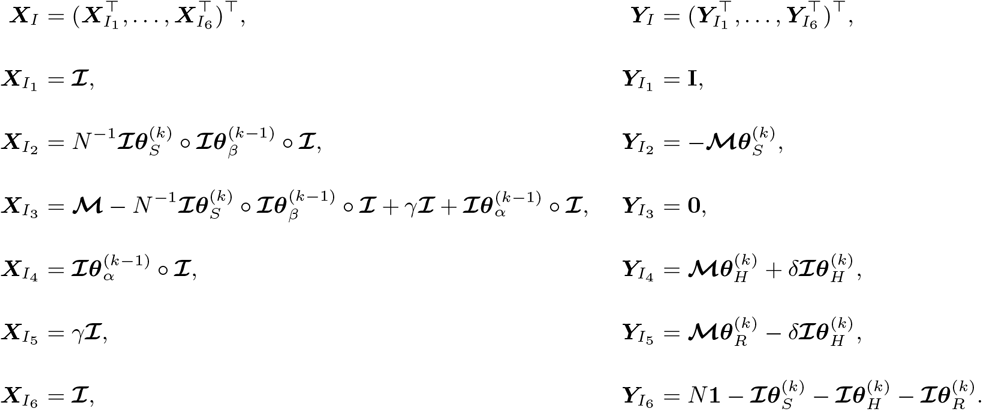

The update for ***θ***_*H*_ is given by 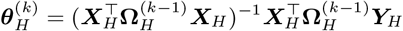, where

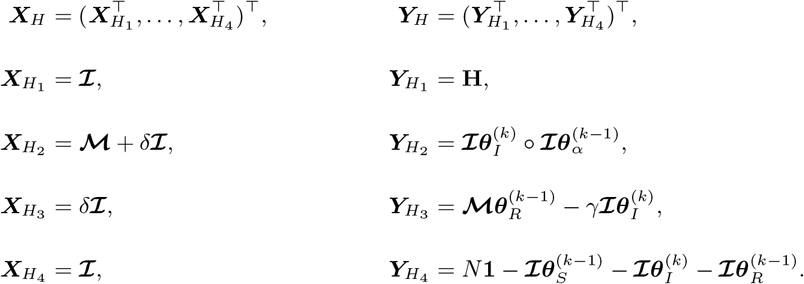

The update for ***θ***_*β*_ is given by 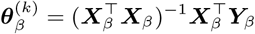, where

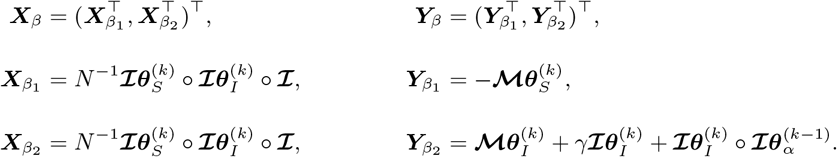

The update for ***θ***_*α*_ is given by 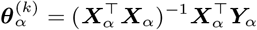, where

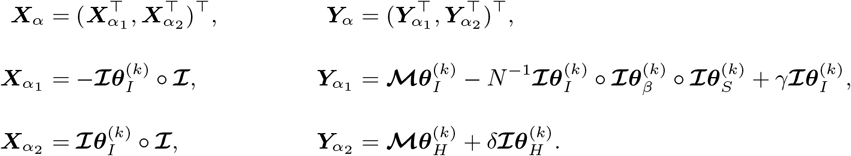

The weight matrices **Ω**_*I*_ and **Ω**_*H*_ are updated at the end of each iteration using the current fitted compartment trajectories, yielding an iteratively reweighted least squares strategy induced by the assumed mean–variance relationships Var(*I*_*t*_) = *ϕ*_*I*_ *I*(*t*) and Var(*H*_*t*_) = *ϕ*_*H*_ *H*(*t*). The weight matrices are thus diagonal, with updates given by

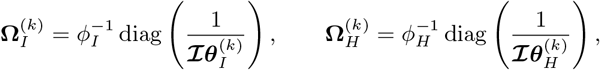

Note that the dispersion parameters cancel in the weighted least squares updates for ***θ***_*I*_ and ***θ***_*H*_, and therefore need not be estimated until the model fitting procedure has converged. Once convergence is achieved, we estimate the dispersion parameters using the traditional method-of-moments estimators with finite-sample corrections. Specifically,

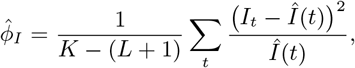

with an analogous estimator used for *ϕ*_*H*_ . Here, *K* denotes the number of observations in **T**_*D*_, and *L* + 1 denotes the number of spline coefficients used in the corresponding compartment representation. Due to the inclusion of the physics-based penalty, the effective degrees of freedom of for the fitted model are bounded by *L*+1. Consequently, the above estimator may slightly overestimate the dispersion parameter in limited data scenarios.

When **T**_*D*_ ≠ **T**_*C*_ and spline representations employ different knot sequences, block updates still follow the same routine outlined above. All design matrices appearing in the physics loss have the same number of rows corresponding to the chosen collocation points, but may have different numbers of columns depending on the knot sequence. For the infectious compartment and its derivative we have **ℐ**_*C,I*_ ***θ***_*I*_ and **ℳ**_*C,I*_ ***θ***_*I*_, where

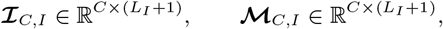

with

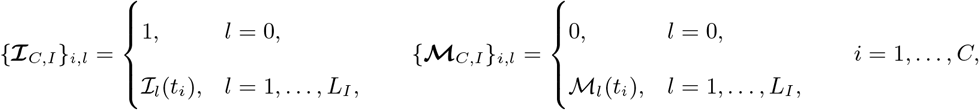

where *C* denotes the number of collocation points, *L*_*I*_ the number of spline basis functions excluding the intercept, and *t*_*i*_ the *i*th collocation point. The corresponding design matrix appearing in the data loss is

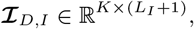

having the same columns but evaluated at the observation times. The spline representations for the remaining compartments and their derivatives are defined analogously.

### A.2 Additional Details on Bootstrapping

Utilizing the detailed bootstrap approach allows for uncertainty quantification of both the compartment trajectories and the time-varying parameters. It also enables uncertainty quantification for smooth functionals of these quantities, such as the effective reproduction number *R*_*t*_(*t*). After refitting the model to each bootstrap data set, we obtain bootstrap estimates of the compartment trajectories and parameters, denoted by 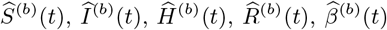, and 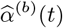. Recalling the definition of *R*_*t*_(*t*), its bootstrap analogue is given by

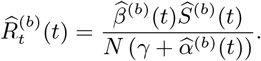

The collection 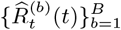 approximates the sampling distribution of *R*_*t*_(*t*), and is used for inference via empirical quantiles.

## Notes

### Competing Interest Statement

The authors have declared no competing interest.

### Funding

- CDC: NU38FT000011

